# Transplantation reveals birthdate-dependent post-mitotic competence in cortical neurons

**DOI:** 10.64898/2026.09.15.751763

**Authors:** Eline Balavoine, Sabine Fièvre, Sergi Roig-Puiggros, Clothilde Ferreira, Julien Prados, Riccardo Bocchi, Denis Jabaudon

## Abstract

Cortical glutamatergic neurons are generated through temporally ordered developmental programs that link neuronal birthdate to laminar position and identity. Yet, because differentiation continues after cell-cycle exit, it remains unclear how far post-mitotic identity can be reshaped by the environment and whether this residual competence varies across neuronal types. Here, we used transplantation to uncouple birthdate from final laminar position by placing newborn neurons in ectopic laminae in the postnatal mouse cortex. E13-born (*i.e.* deep layer-destined) and E15-born (*i.e.* superficial layer-destined) donor neurons displayed distinct birthdate-associated molecular programs before transplantation. After grafting into P0 hosts, both populations migrated and settled across cortical layers, enabling comparison across laminar environments. We simultaneously profiled transcriptomic, morphological and electrophysiological neuronal identities using Patch-seq. Transplanted neurons developed pyramidal morphologies and active membrane properties, but their final identity was not imposed by their laminar position. Thus, the postnatal cortex supports integration and maturation but does not impose canonical laminar identity. Critically, neuronal competence was asymmetric across birthdates. E15-born neurons remained committed toward a robust superficial layer neuron-like profile independently of cortical position, whereas E13-born neurons retained a broad, non-canonical differentiation rather than either converging on a deep- or superficial-layer reference state. This is consistent with the wider intrinsic repertoire of early-born neurons, which the postnatal environment does not resolve. Thus, post-mitotic identity is constrained by developmental history in a birthdate-dependent manner, with early- and late-born neurons retaining fundamentally different differentiation potentials after cell-cycle exit.

## Introduction

The mammalian neocortex contains diverse classes of glutamatergic neurons that differ in laminar position, molecular identity, dendritic architecture, electrophysiological properties and connectivity^1–6^. These identities emerge through a temporally ordered neurogenic program, during which cortical progenitors sequentially generate these distinct neuronal classes in a laminar inside-out sequence^7–12^. Early-born neurons preferentially populate deep cortical layers and include extratelencephalic (ET) and intratelencephalic (IT) projection neuron classes, whereas later-born neurons settle in superficial layers and are mainly associated with IT identities^7,8,10,11,13–15^. Thus, neuronal birthdate is tightly linked to neuronal subtype specification and final laminar organization across mammals^7–15^.

Although cortical glutamatergic neuron identities are initiated during embryonic neurogenesis, their differentiation continues after cell-cycle exit^12,16–18^. Newly generated neurons migrate through the developing cortex, settle into appropriate laminar positions and progressively acquire mature morphological, electrophysiological and transcriptional features^12,16–19^. This extended period of post-mitotic differentiation raises an important question: to what extent are neuronal identities determined by birthdate-dependent developmental programs, and to what extent can they be modified by post-mitotic environment? This question is particularly relevant to distinguishing cell-intrinsic constraints from extrinsic influences associated with cortical laminar position and developmental timing^9,20–23^, and to determining whether post-mitotic responsiveness to environmental cues varies across birthdates.

Transplantation provides a powerful approach to untangle these contributions^9,21–23^. By placing embryonic cortical neurons in different locations of a host cortex, stage-specific donor neurons of defined birthdate can be exposed to a different laminar environment by occupying positions that are either consistent with, or heterotopic to, their normal laminar fate. This experimental design uncouples donor birthdate from final radial position^9,21–23^, enabling assessment of the relative contributions of intrinsic birthdate-associated programs and extrinsic laminar environment to neuronal morphology, intrinsic electrophysiological properties and transcriptional identity^24–26^.

Here, we combined FlashTag (FT) birthdating^16,27^, transplantation^21,28^ and Patch-seq^24^ to examine how embryonic cortical neurons differentiate after transplantation across laminar positions in the early postnatal mouse cortex. We labelled E13- and E15-born donor neurons, corresponding mainly to early-born L5-fated and late-born L2/3-fated populations, respectively, and transplanted them into P0 host cortex, providing a common postnatal environment in which deep and superficial layers are becoming discernible and extrauterine influences have begun. We then compared the post-transplantation differentiation of donor neurons across layers. By integrating single-cell laminar position, morphology, intrinsic electrophysiology and transcriptional identity, we tested how final neuronal identity reflects the host environment versus neuronal birthdate, and whether this relationship differs between early- and late-born neurons. Our results show that transplanted neurons enter the postnatal cortical plate and integrate into both superficial and deep layers, but their subsequent differentiation differs according to birthdate. E15-born neurons remain consistently committed toward a L2/3-like trajectory, whereas E13-born neurons display broader, mixed identity outcomes rather than uniformly acquiring a L5 fate. Together, these findings reveal birthdate-dependent post-mitotic competence: the postnatal cortex supports neuronal integration and maturation but only partially redirects neuronal identity. Late-born neurons remain canalized toward a L2/3 trajectory, whereas early-born neurons retain a broader competence that the postnatal environment does not consolidate into a single canonical fate.

## Results

### Donor neurons have birthdate-associated molecular identities

During corticogenesis, neuronal birthdate is tightly linked to the acquisition of distinct laminar and projection identities^7,10,11,29^. Glutamatergic neurons born around E13 typically settle in deep cortical layers and give rise to both ET and IT projection neurons, whereas neurons born around E15 preferentially populate superficial layers and are predominantly IT (Fig. 1a, right). We therefore labelled embryos at E13.5 or E15.5 with FT (hereafter and throughout the manuscript E13 and E15 for simplicity, with E0.5 defined as the day of plug detection) and examined the labelled cohorts at P7. This strategy generated two spatially segregated populations, with E13-labelled neurons located predominantly in deep layers and E15-labelled neurons in superficial layers (Fig. 1a, left).

**Fig. 1.**
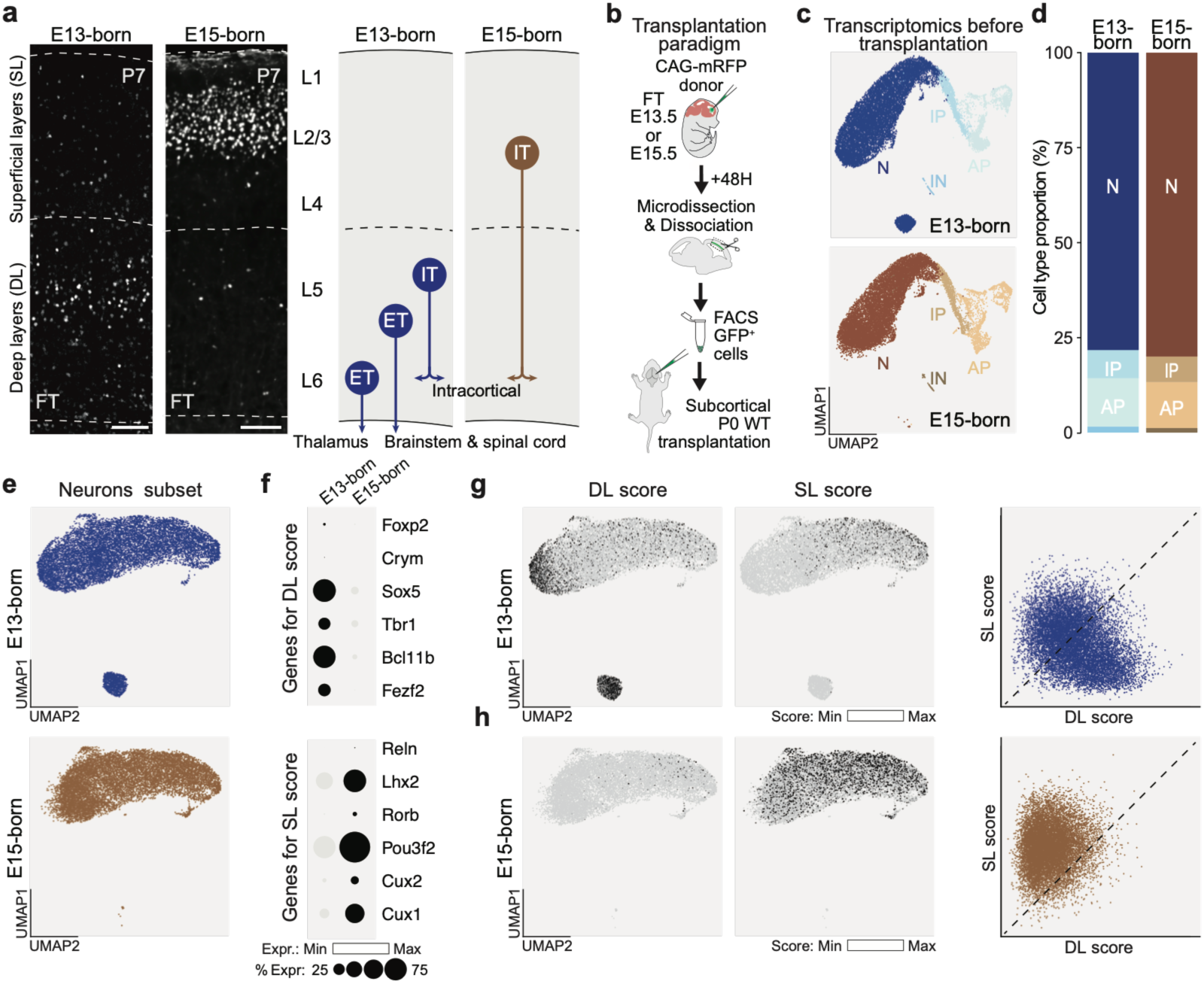
Donor neurons show birthdate-associated molecular identities before transplantation. **(a)** Representative coronal sections of FT-labelled brains injected at E13 or E15 and collected at P7, together with a schematic summary of the expected laminar positions and projection identities of each birthdate-labelled population. E13-born neurons mainly populate DL and include extratelencephalic (ET) and intratelencephalic (IT) projection neurons, whereas E15-born neurons populate SL and are mainly IT neurons. **(b)** Heterochronic transplantation workflow. CAG-mRFP donor embryos were labelled with FT at E13 or E15, collected 48 h later, microdissected from presumptive S1, dissociated, FACS-purified, and transplanted into P0 wild-type hosts. **(c)** UMAP representation of donor-cell scRNA-seq profiles before transplantation. N, neurons; IP, intermediate progenitors; AP, apical progenitors; IN, interneurons. **(d)** Cell-type proportions in E13-born and E15-born donor preparations, showing enrichment for post-mitotic neurons after FT labelling and FACS purification. **(e)** UMAP representation of the post-mitotic neuronal subset used for downstream analyses. **(f)** Dot plot showing expression of canonical DL and SL marker genes. Dot size indicates the percentage of expressing cells, and color intensity indicates normalized expression. **(g, h)** DL and SL module scores in E13-born (g) and E15-born (h) neurons. Scale bars, 100 μm.

We next used single-cell RNA sequencing (scRNA-seq) to define the cellular and molecular state of E13- and E15-born neurons at the time of transplantation, before exposure to the postnatal host environment. CAG-mRFP donor embryos were labelled with FT at E13 or E15, collected 48 h later, and the presumptive somatosensory (S1) cortex was microdissected, dissociated and FACS-purified for scRNA-seq analysis (Fig. 1b). Both purified donor preparations were strongly enriched for post-mitotic neurons (>75%), with limited fractions of intermediate progenitors, apical progenitors and other cell classes (Fig. 1c, d; Supplementary Fig. 1a). To determine the fraction of FT-positive donor cells that were proliferative at the time of transplantation, we used two complementary EdU-labelling paradigms. In the first paradigm, EdU was administered as a short pulse 47 h after FT labelling, 1 h before donor collection, to identify cells that were in S-phase immediately before grafting (Supplementary Fig. 1b, top). In the second paradigm, EdU was provided chronically during the 24 h preceding donor collection, to capture cells that had entered S-phase over a broader pre-transplantation window (Supplementary Fig. 1b, bottom). In both conditions, EdU incorporation was detected in only a very small fraction of FT-positive cells, indicating that most labelled donor cells had exited the cell cycle and were largely non-proliferative at the time of transplantation (Supplementary Fig. 1c).

We then asked whether birthdate-associated molecular identities were already detectable in these newly generated donor neurons before transplantation. Restricting the analysis to the post-mitotic neuronal subset revealed birthdate-specific transcriptional signatures between E13-born and E15-born neurons (Fig. 1e). E13-born neurons preferentially expressed deep-layer-associated genes (Fig. 1f, top; Supplementary Fig. 1d, e), including markers linked to layer 5/6 neuron identity, and displayed higher deep-layer module scores (Fig. 1g). Conversely, E15-born neurons were enriched for superficial-layer-associated programs (Fig. 1f, bottom; Supplementary Fig. 1d, f), including Cux-family genes and Unc5d-related signatures, and showed higher superficial-layer module scores (Fig. 1h). Hence, birthdate-associated programs are already established in newly post-mitotic donor neurons, providing a defined molecular starting point against which post-transplantation differentiation can be measured.

To ensure that these transcriptional differences reflected birthdate identity rather than differences in maturation state, we further performed pseudomaturation analysis using a published developmental reference^12^. Both E13- and E15-born donor neurons, collected 48 h after FT labelling, were positioned between the published 24 h and 96 h post-labelling states and showed comparable maturation progression (Supplementary Fig. 1g). Together, these analyses indicate that both donor preparations consisted largely of newly post-mitotic neurons and showed broadly comparable maturation progression, despite displaying distinct birthdate-associated molecular identities before exposure to the host environment.

### Transplanted neurons integrate throughout the host postnatal cortical column

At P0, the neocortex is still immature and continues to support the migration of superficial-layer neurons^30,31^. We therefore asked whether embryonic donor neurons could migrate within the host cortical plate and whether their radial distribution would remain constrained by donor birthdate. This approach exposes E13- and E15-born neurons to the same postnatal environment while dissociating embryonic birthdate from final laminar position. Following transplantation close to the periventricular white matter of P0 (Fig. 1b), both E13-born and E15-born RFP-positive donor cells were initially detected around the transplantation site. Over the first postnatal week, these cells progressively redistributed into the host cortex (Fig. 2a, b). Quantification across P1.5, P3.5 and P5.5 showed that both donor populations shifted away from periventricular and subcortical positions, with most RFP-positive cells entering the cortical plate by P5.5 (Supplementary Fig. 2a, b). Thus, unlike adult cortical transplantation paradigms in which grafted neurons mature and integrate within a relatively restricted host territory^32^, the early postnatal cortex allowed embryonic donor neurons to disperse radially into the cortical plate.

**Fig. 2.**
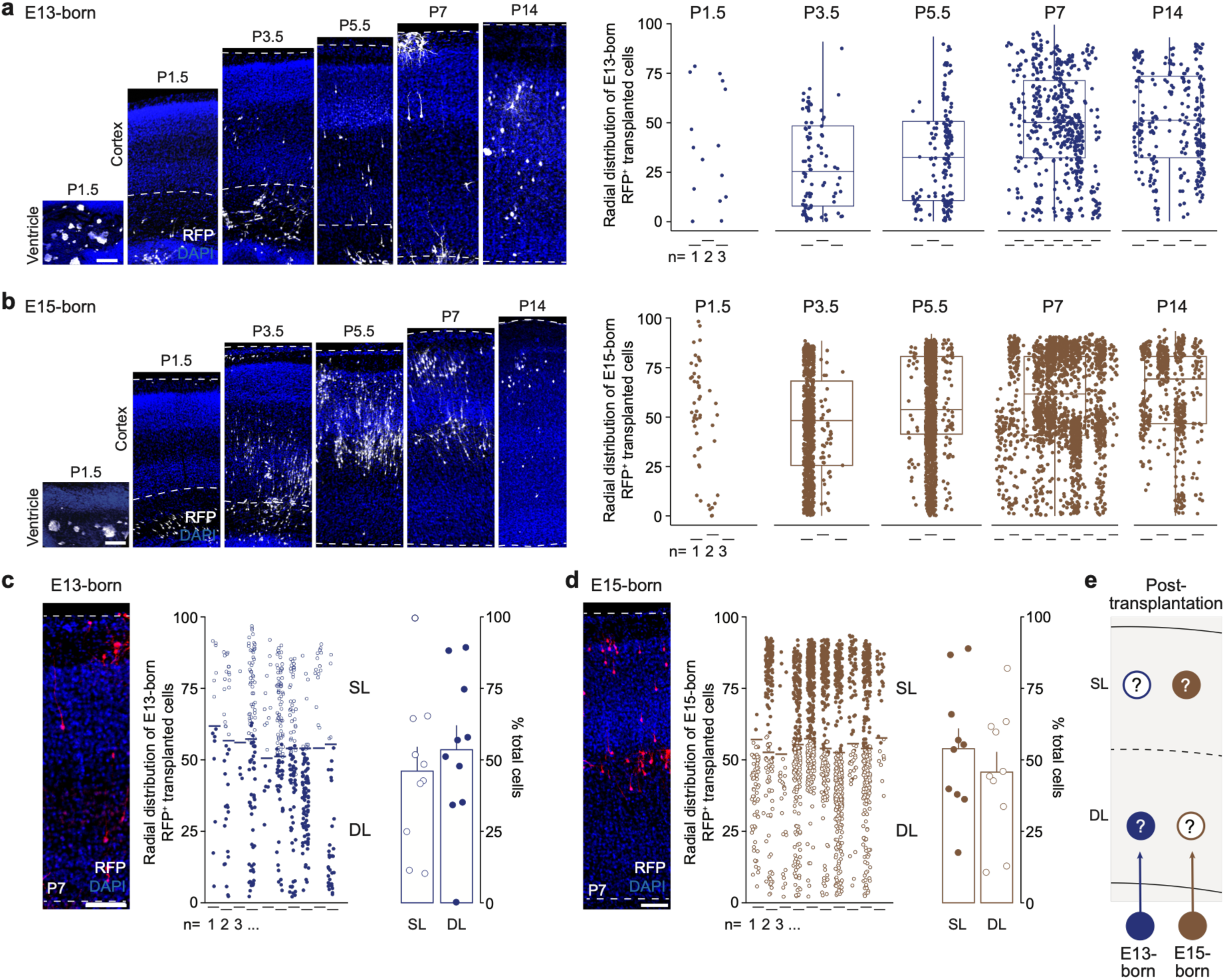
Transplanted embryonic neurons migrate into superficial and deep cortical layers. **(a, b)** Representative coronal sections showing RFP-labelled E13-born (a) and E15-born (b) transplanted cells collected at P1.5, P3.5, P5.5, P7, and P14. Quantification on the right shows the radial distribution of individual integrated E13-born (a) and E15-born (b) neurons across postnatal time points. Cell position was normalized to cortical thickness in each slice, with 0 corresponding to the bottom of L6 and 100 to the top of L1. Circles represent individual transplanted neurons; ticks below the x-axis indicate biological replicates/transplanted pups. **(c, d)** Representative P7 sections and radial distribution of E13-born (c) and E15-born (d) neurons assigned to SL or DL positions. Filled circles indicate neurons located in the expected birthdate-associated laminar compartment, whereas open circles indicate heterotopic positions. n = 10 transplanted mice. **(e)** Schematic showing that both donor populations integrate into SL and DL positions after transplantation, enabling comparison between birthdate-encoded identity and laminar environmental influence. Values are shown as mean +/− SEM. Scale bars, 100 μm.

We then examined whether donor birthdate influenced the radial position reached by transplanted neurons. At P7, RFP-positive donor cells were broadly distributed across the radial axis of the host cortex, with both E13-born and E15-born neurons recovered in superficial as well as deep cortical layers (Fig. 2c, d). Thus, E13-born neurons, which normally preferentially populate deep layers (homotopic), could also be found in superficial positions (heterotopic), whereas E15-born neurons, which normally populate superficial layers (homotopic), could also be recovered in deep positions (heterotopic).

We next verified that this broad cortical distribution reflected integration of post-mitotic donor neurons rather than local expansion of rare proliferative cells present at transplantation. Using the same EdU-labelling paradigms described above, we analyzed transplanted cells at P7, after 7 days in the host cortex. In both paradigms, integrated RFP-positive donor cells showed no EdU incorporation, indicating that the integrated neurons were not derived from proliferative donor cells present at the time of transplantation (Supplementary Fig. 2c, d). In addition, most donor cells expressed the mature neuronal marker NeuN and showed no co-localization with the astrocytic marker Sox9, confirming that the analyzed donor population was predominantly neuronal (Supplementary Fig. 2e, f).

Together, these results show that embryonic donor neurons can migrate into the postnatal cortical plate and occupy both superficial and deep radial positions, providing a paradigm in which neuronal birthdate is uncoupled from final radial position. This transplantation paradigm therefore provides a framework to compare the contribution of birthdate-associated identity with that of the postnatal laminar environment (Fig. 2e).

### Transplanted neurons acquire pyramidal morphologies with birthdate-associated features

Having established that transplanted neurons can occupy both homotopic and heterotopic positions in the postnatal cortex, we next asked how differentiation was associated with donor birthdate and final laminar position. To simultaneously interrogate single-neuron morphology, electrophysiology and transcriptomics, we performed Patch-seq^24^ of RFP-positive transplanted neurons and neighboring RFP-negative host neurons on acute cortical slices at P14-P16. Following whole-cell recording, neurons were filled with biocytin for morphological reconstruction and their nuclei were collected for scRNA-seq (Fig. 3a). The final Patch-seq dataset included neighboring control L2/3 and L5 neurons, as well as E13-born and E15-born transplanted neurons located in either L2/3 or L5 (Fig. 3b).

**Fig. 3.**
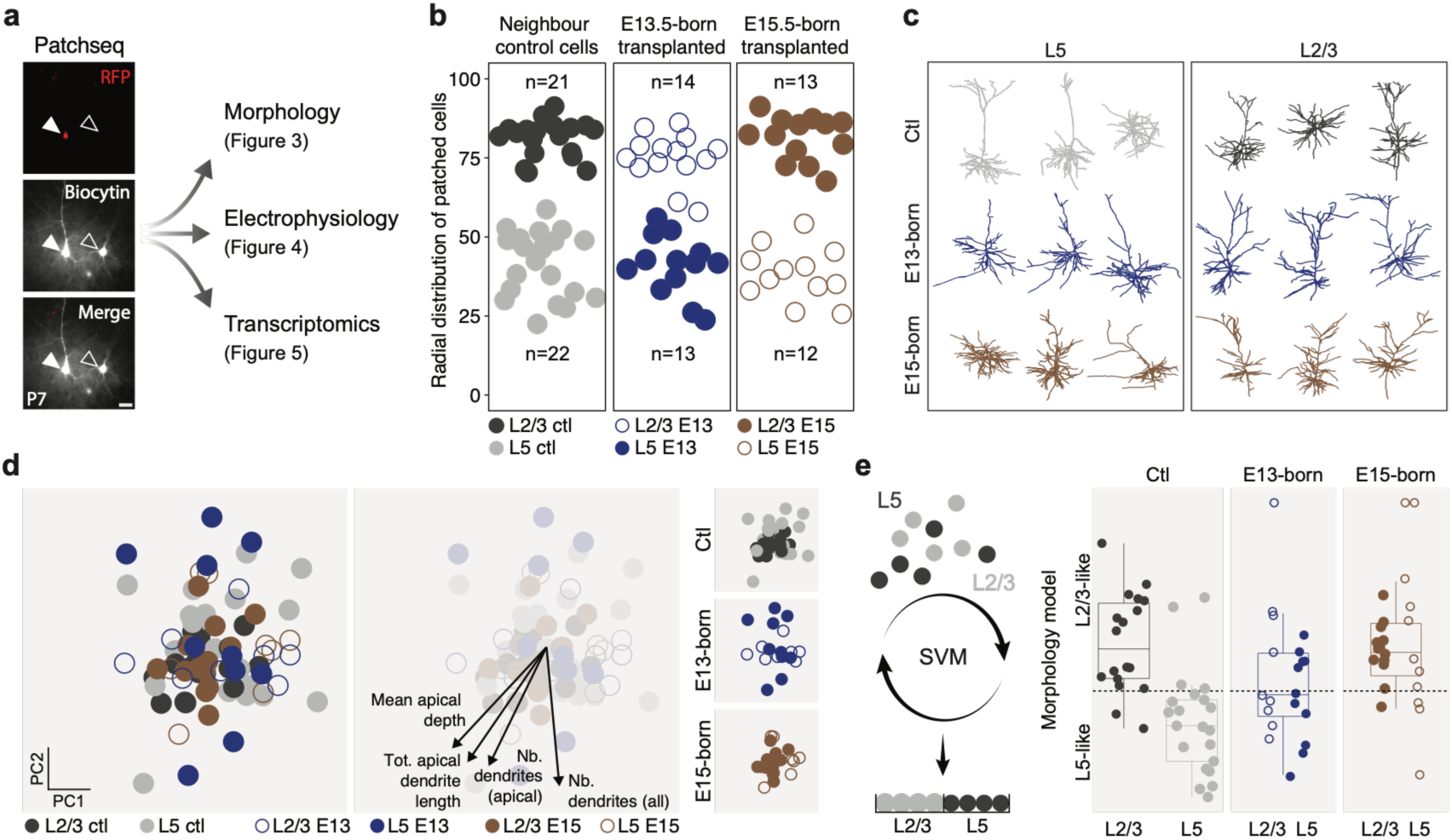
Morphological analysis reveals heterogeneous identities in transplanted neurons. **(a)** Patch-seq workflow used to combine morphological, electrophysiological, and transcriptomic analyses from the same neurons. RFP-positive transplanted neurons were targeted in acute slices, filled with biocytin during whole-cell recording, reconstructed, and processed for RNA-seq. **(b)** Radial distribution of patched control neurons and transplanted E13-born and E15-born neurons used for morphological, electrophysiological, and transcriptomic analyses in Figs. 3-5. Cells were grouped according to their final position in L2/3 or L5. **(c)** Representative reconstructed morphologies of control, E13-born, and E15-born neurons located in L5 or L2/3. **(d)** PCA of morphological features from patched control and transplanted neurons. The four most discriminative variables are highlighted on the right. **(e)** SVM classifier trained and cross-validated on control L2/3 and L5 neurons using morphological features, then applied to transplanted neurons to assign each cell a continuous L2/3-like to L5-like score. The grey line marks the L2/3-like/L5-like classification threshold. n = 21 neurons for L2/3 ctl, 22 for L5 ctl, 14 for L2/3 E13, 13 for L5 E13, 13 for L2/3 E15, and 12 for L5 E15. Values are shown as mean +/− SEM. Scale bar, 15 μm.

Biocytin reconstructions showed that transplanted neurons developed pyramidal-like morphologies, with apical and basal dendritic arbors (Fig. 3c; Supplementary Fig. 3a, left). We used 54 quantitative morphological parameters to compare reconstructed control and transplanted neurons (Supplementary Fig. 4a). Principal component analysis (PCA) of these values revealed that neighboring control L2/3 and L5 neurons occupied distinct regions of the morphological PCA space, capturing layer-associated differences among endogenous cortical glutamatergic neurons (Fig. 3d).

We then asked whether transplanted neurons were morphologically closer to neurons of their final laminar position or instead retained birthdate-associated features. Hierarchical analysis revealed that transplanted neurons did not segregate simply according to their final laminar position (Supplementary Fig. 3a, right). To quantify this more directly, we trained a support vector machine classifier (SVM) on neighboring control L2/3 and L5 neurons and applied it to transplanted cells. E13-born transplanted neurons showed heterogeneous morphology scores, with both L2/3-like and L5-like features, regardless of whether they occupied homotopic or heterotopic positions. In contrast, E15-born neurons were more consistently classified as L2/3-like, including when recovered from L5 (Fig. 3e).

Together, these results show that transplanted embryonic neurons mature into pyramidal-like cells within the postnatal cortex, but that their morphological differentiation is not simply dictated by the layer in which they settle. Instead, E15-born neurons were consistently biased toward a superficial-layer-like morphology, whereas E13-born neurons showed broader variability and mixed L2/3-like and L5-like features rather than a uniform deep-layer morphology.

### Transplanted neurons show L2/3-biased electrophysiological properties

We next asked whether transplanted neurons acquired electrophysiological properties corresponding to their donor birthdate or to their final laminar position. Using the Patch-seq recordings from the same dataset, we first examined representative current-clamp responses and individual intrinsic electrophysiological parameters. Control L2/3 and L5 neurons displayed distinct firing behaviors, whereas transplanted neurons generally showed regular spiking activity. E13-born neurons showed greater electrophysiological variability, while E15-born neurons displayed more homogeneous responses resembling those of L2/3 controls (Fig. 4a). This greater variability among E13-born neurons mirrors the morphological heterogeneity described above.

**Fig. 4.**
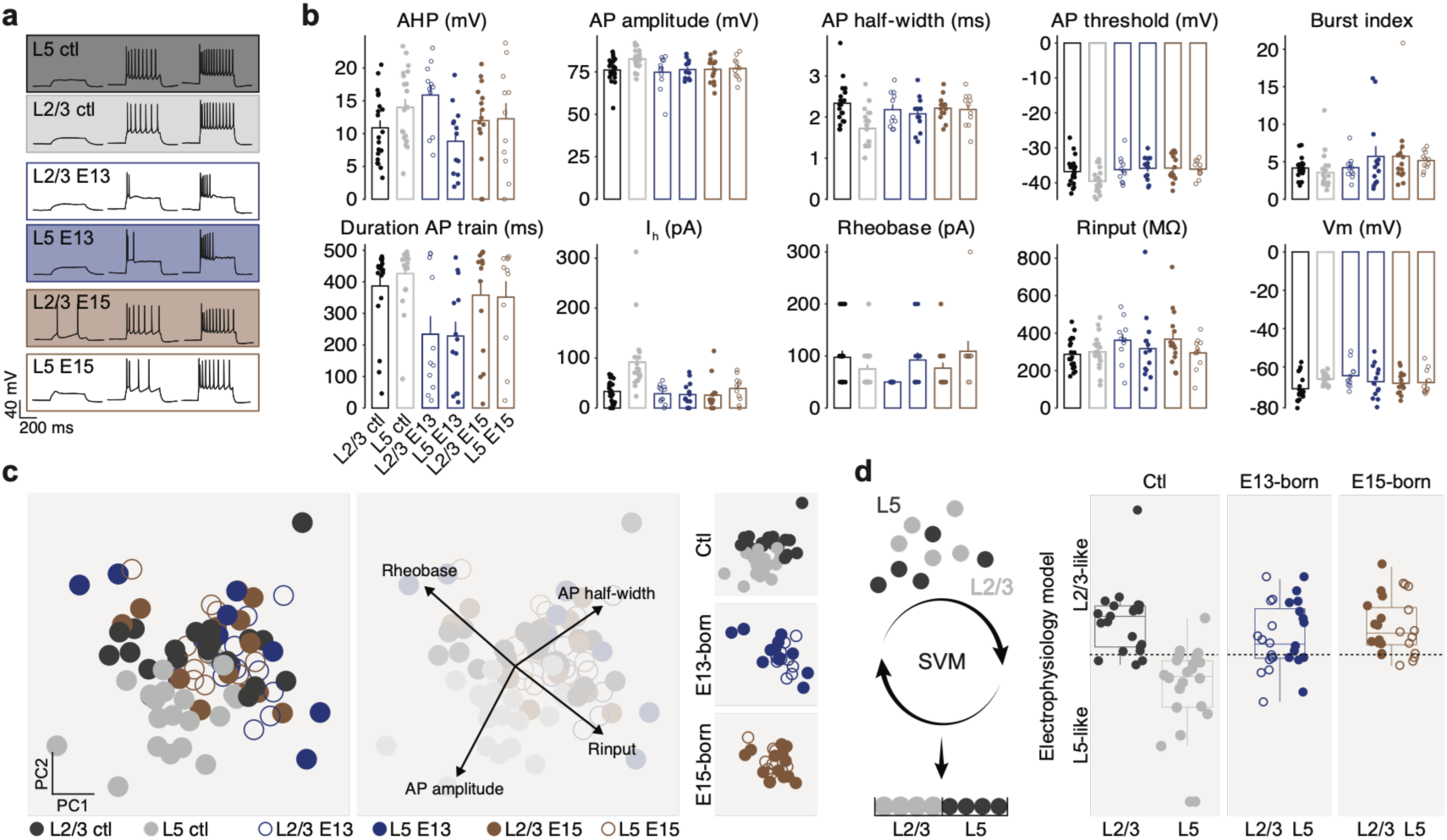
Transplanted neurons are electrophysiologically biased toward L2/3-like properties. **(a)** Representative current-clamp traces from control L5, control L2/3, E13-born L2/3-positioned, E13-born L5-positioned, E15-born L2/3-positioned, and E15-born L5-positioned neurons. **(b)** Quantification of electrophysiological parameters across groups. n = 21 neurons for L2/3 ctl, 22 for L5 ctl, 14 for L2/3 E13, 13 for L5 E13, 13 for L2/3 E15, and 12 for L5 E15. **(c)** PCA of electrophysiological features measured from patched control and transplanted neurons. The four most discriminative variables are highlighted on the right. **(d)** SVM classifier trained and cross-validated on control L2/3 and L5 neurons using electrophysiological features, then applied to transplanted neurons to assign each cell a continuous L2/3-like to L5-like score. The grey line marks the L2/3-like/L5-like classification threshold. Values are shown as mean +/− SEM.

Parameter-level analyses further showed that L5-like electrophysiological features were not robustly acquired by transplanted neurons, even when these cells integrated into L5. Features that contributed to the distinction between control L2/3 and L5 neurons, including hyperpolarization-activated current-related properties, rheobase, action-potential waveform measures and firing-train parameters, remained overall closer to L2/3 values in the transplanted populations (Fig. 4b).

We then used multivariate analyses to integrate these electrophysiological features across neighboring host neurons and transplanted cells. PCA separated control L2/3 and L5 neurons, indicating that the recorded parameters captured layer-associated electrophysiological differences among endogenous cortical glutamatergic neurons (Fig. 4c). In contrast, transplanted neurons clustered closer to L2/3 controls across donor birthdate and final laminar position.

To quantify this relationship, we trained an electrophysiology-based SVM classifier on neighboring control L2/3 and L5 neurons and applied it to transplanted cells. Most transplanted neurons were classified as L2/3-like, including cells recovered from L5 positions (Fig. 4d). Together, these results indicate that electrophysiological differentiation after transplantation was globally biased toward L2/3-like properties and showed limited evidence for layer-instructed acquisition of a canonical deep-layer functional profile. Thus, electrophysiology points to a permissive but L2/3-biased postnatal environment, within which E13-born neurons retained greater variability than the more homogeneous E15-born population.

### Patch-seq reveals asymmetric transcriptomic differentiation after transplantation

We next asked how the molecular identity of transplanted neurons was associated with donor birthdate and final laminar position. Using the Patch-seq transcriptomes from the same cells analyzed morphologically (Fig. 3) and electrophysiologically (Fig. 4), we first performed PCA to compare neighboring host neurons and transplanted cells. Control L2/3 and L5 neurons separated along this transcriptomic PCA space, whereas transplanted neurons clustered closer to one another than to either control group (Fig. 5a). Pairwise correlation analysis supported this observation, with transplanted neurons showing higher similarity across donor birthdates and final positions than expected from a simple layer-matching model (Fig. 5b). These results suggest that transplanted neurons adopted a shared molecular state that was not determined solely by their final laminar position.

**Fig. 5.**
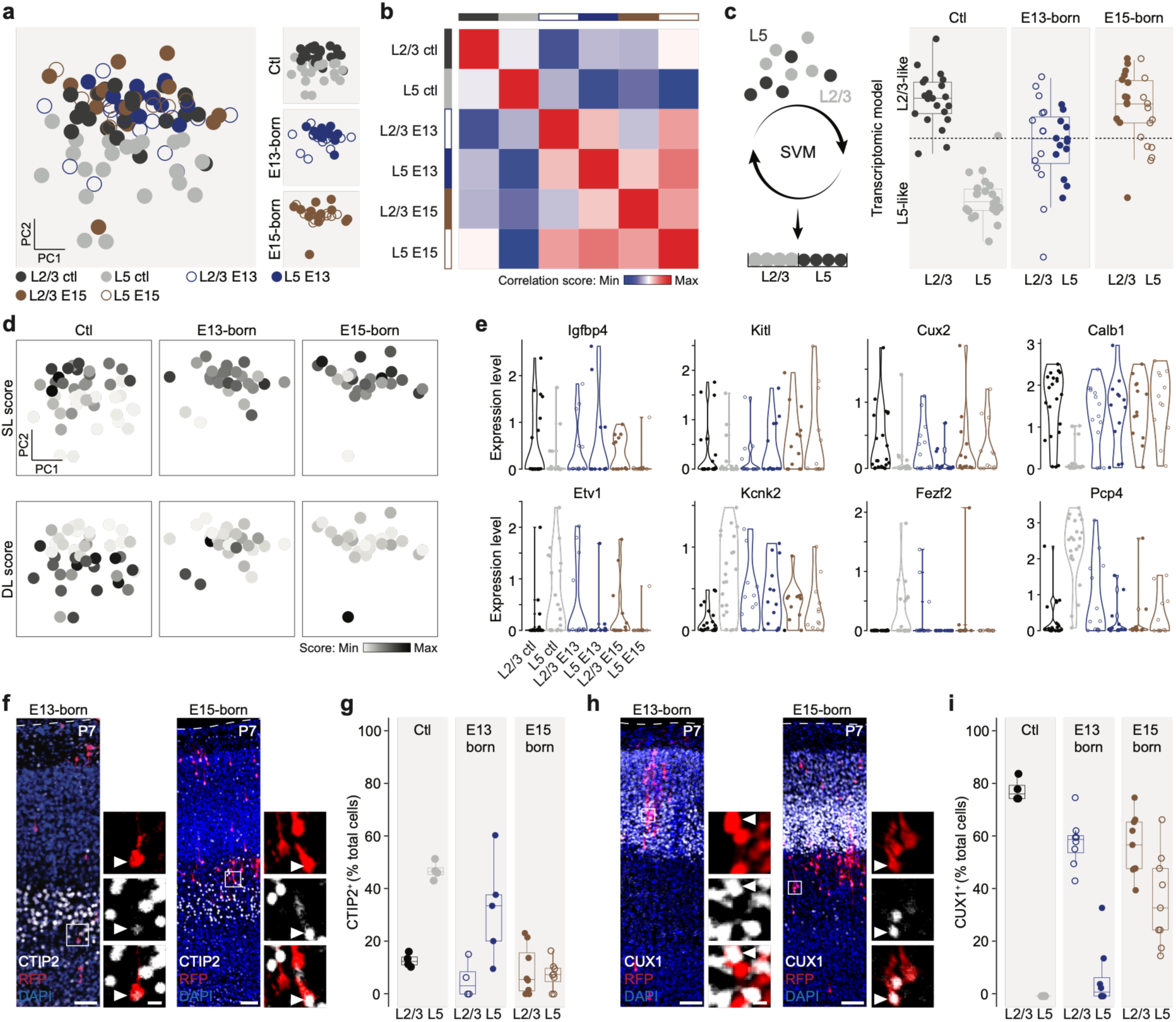
Patch-seq transcriptomes reveal asymmetric differentiation with a stronger L2/3-like bias in E15-born neurons. **(a)** PCA of transcriptomic features measured from patched control and transplanted neurons. **(b)** Correlation matrix based on pairwise correlations between group means in reduced PCA space, showing that transplanted neurons are more similar to one another than to either control group. **(c)** SVM classifier trained and cross-validated on control L2/3 and L5 neurons using transcriptomic features, then applied to transplanted neurons to assign each cell a continuous L2/3-like to L5-like score. The grey line marks the L2/3-like/L5-like classification threshold. **(d)** SL and DL module scores projected onto transcriptomic PCA space. **(e)** Violin plots showing single-cell expression of selected L2/3- and L5-associated genes. **(f-i)** Representative CTIP2 (f, g) and CUX1 (h, i) immunostaining and quantification in RFP-positive E13-born and E15-born transplanted neurons at P7, compared with control L2/3 and L5 neurons. Arrowheads indicate marker-positive transplanted cells. In (a-e), n = 21 neurons for L2/3 ctl, 22 for L5 ctl, 14 for L2/3 E13, 13 for L5 E13, 13 for L2/3 E15, and 12 for L5 E15. In (f-i), n = 4 control mice, 5 E13-born transplanted mice, and 9 E15-born transplanted mice. Values are shown as mean +/− SEM. Scale bars, 100 μm; insets, 15 μm.

We then assessed whether transplanted neurons retained or acquired layer-associated molecular programs. A transcriptome-based SVM classifier trained on neighboring control L2/3 and L5 neurons classified E15-born transplanted neurons predominantly as L2/3-like, including when they were recovered from L5 positions. In contrast, E13-born neurons showed more heterogeneous scores, with mixed L2/3-like and L5-like features and less consistent assignment to either class (Fig. 5c). Module-score analysis further supported this distinction: L2/3-associated programs were broadly detected in transplanted neurons, whereas deep-layer-associated programs were more variable and most evident in subsets of E13-born cells (Fig. 5d).

Single-gene analysis provided additional resolution of these mixed identities. Transplanted neurons expressed several genes associated with superficial-layer/IT programs, while E13-born neurons also showed partial expression of genes associated with deep-layer identities (Fig. 5e). Thus, the transcriptomic data point to a shared post-transplantation state, superimposed on molecular birthdate-dependent bias: E15-born neurons were more consistently L2/3-like, whereas E13-born neurons retained broader and more mixed identity features.

We next examined whether these transcriptomic trends were reflected at the protein level. At P7, E15-born transplanted neurons frequently expressed CUX1 and showed little CTIP2 expression, even when located in deeper cortical positions. In contrast, E13-born transplanted neurons displayed more variable marker expression, including CUX1 expression in superficial positions and incomplete or heterogeneous CTIP2 expression in deep positions (Fig. 5f-i). Together, these transcriptomic and immunohistochemical analyses argue against a simple model in which the host layer fully instructs transplanted neuron identity. Instead, they reveal asymmetric post-mitotic differentiation: E15-born neurons retain a stable superficial/IT-like molecular bias, whereas E13-born neurons occupy a broader, less canonical molecular space, consistent with the diverse intrinsic repertoire of early-born glutamatergic neurons represented in the donor population rather than with the acquisition of a stable L5 identity imposed by the deep-layer environment.

### Multimodal integration confirms asymmetric birthdate-specific differentiation

Finally, we asked whether integrating morphology, electrophysiology and transcriptomics could better resolve the identity of transplanted neurons than single modalities. To address this, we generated a multimodal identity model trained on neighboring control L2/3 and L5 neurons and then applied it to transplanted cells. Each individual modality discriminated control laminar identities with high accuracy, and combining all three modalities further improved classification performance (Fig. 6a, b).

**Fig. 6.**
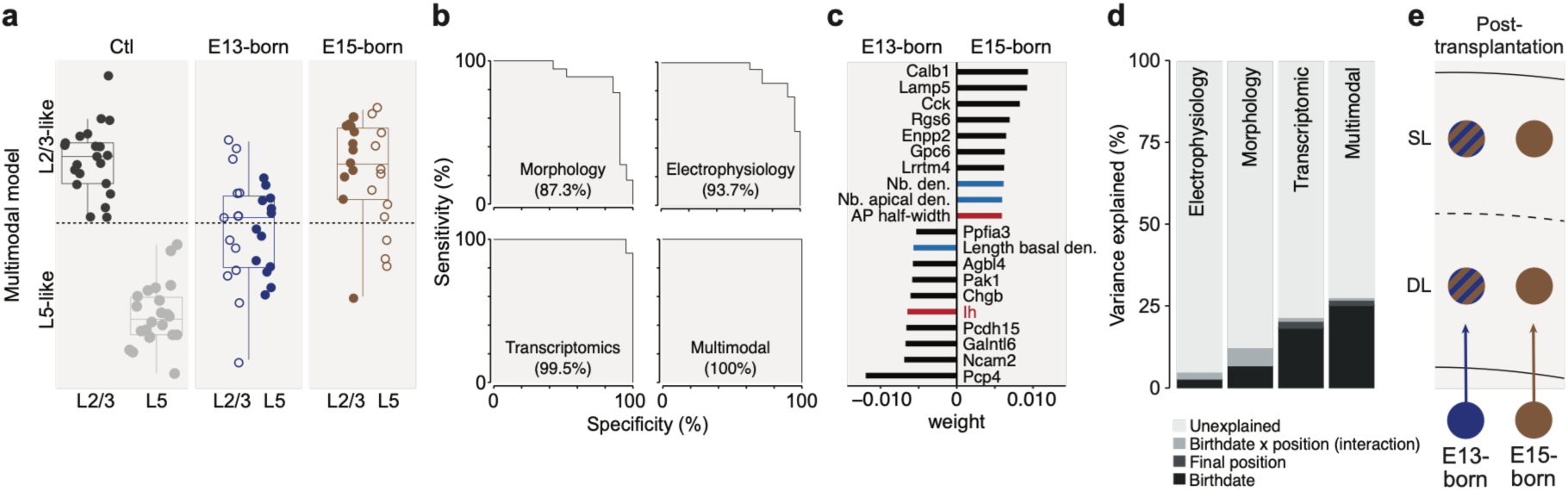
Multimodal integration identifies asymmetric birthdate-specific differentiation after transplantation. **(a)** Multimodal SVM classifier trained and cross-validated on control L2/3 and L5 neurons using combined morphological, electrophysiological, and transcriptomic features, then applied to transplanted neurons to assign each cell a continuous L2/3-like to L5-like score. The grey line marks the L2/3-like/L5-like classification threshold. **(b)** ROC curves for morphology, electrophysiology, transcriptomics, and the integrated multimodal model. Values indicate model performance for distinguishing L2/3-like and L5-like identities in the control training set. **(c)** Top weighted features contributing to multimodal classification. Black bars indicate transcriptomic features, blue bars indicate morphological features, and red bars indicate electrophysiological features. **(d)** Variance partitioning among transplanted neurons across morphology, electrophysiology, transcriptomics, and multimodal identity, showing the relative contribution of birthdate, final position, their interaction, and unexplained variance. **(e)** Summary model. E15-born neurons retain a stable SL/L2/3-like identity after transplantation, irrespective of final position. E13-born neurons show broader mixed identity outcomes and do not acquire a homogeneous canonical L5-like identity after transplantation.

Applying this integrated model to transplanted neurons revealed clear differences between donor groups. E15-born neurons were predominantly classified as L2/3-like, regardless of their radial position, indicating a consistent commitment toward late-born neuronal properties. In contrast, E13-born neurons showed more heterogeneous multimodal scores and were distributed across the L2/3-like to L5-like axis, indicating that early-born neurons did not converge on a single identity state or uniformly recover a canonical L5-like fate after transplantation (Fig. 6a).

We next examined which features contributed most strongly to the integrated classification. Transcriptomic features ranked prominently among the highest-weighted predictors, but morphological and electrophysiological parameters were also represented among the informative features (Fig. 6c). Thus, laminar identity was not captured by a single readout but instead emerged from concordant information across modalities.

To compare the relative contribution of donor birthdate and final laminar position across modalities, we performed variance partitioning analysis. In morphology, electrophysiology, transcriptomics and the multimodal model, donor birthdate explained more variance than final position, whereas the interaction between birthdate and position contributed less. However, a substantial fraction of variance remained unexplained, consistent with the heterogeneous, non-canonical outcomes of transplanted neurons, particularly within the E13-born population (Fig. 6d).

Together, these analyses support a model of asymmetric post-mitotic differentiation rather than a simple hierarchy in which birthdate fully determines identity. E15-born neurons appear consistently biased toward a superficial-layer/L2/3-like state irrespective of final position, whereas E13-born neurons show broader heterogeneous outcomes but do not resolve into a homogeneous deep-layer/L5-like identity after transplantation (Fig. 6e).

## Discussion

Cortical glutamatergic neuron identities emerge through temporally regulated developmental programs that link neuronal birthdate to laminar position and projection subtype^12,16^. Classical transplantation studies established that cortical neurons respond to extrinsic cues within restricted windows of competence, and that the developmental state of the host environment influences laminar fate and projection identity^23,33,34^. Here, we revisited this question at the post-mitotic stage by combining FT birthdating, heterochronic transplantation and multimodal Patch-seq^24^. We show that E13-born and E15-born donor neurons already carried distinct birthdate-associated molecular programs before transplantation, migrated into both superficial and deep positions of the postnatal host cortex, but did not simply adopt the identity predicted by their final laminar position. Previous developmental studies have shown that cortical neurons can acquire key subtype-associated molecular and connectivity features despite substantial disruption of their normal laminar position^35,36^. Our findings confirm and extend this principle by showing that post-mitotic neurons transplanted into a heterochronic postnatal environment likewise do not simply adopt the identity associated with their final laminar position.

Our findings indicate that the postnatal cortex is permissive for survival, migration and differentiation of embryonic donor neurons, but has limited capacity to impose laminar identity on post-mitotic neurons. Both E13-born and E15-born neurons entered the cortical plate and developed neuronal features, including pyramidal-like morphologies and active electrophysiological properties. However, their differentiation outcomes differed markedly according to donor birthdate.

This birthdate-dependent asymmetry is the central biological feature of the transplantation response. E15-born neurons retained a stable superficial-layer/L2/3-like identity across modalities, even when recovered from deep positions. This supports the idea that later-born neurons become restricted toward an intratelencephalic/superficial-layer trajectory shortly after neurogenesis and are only weakly redirected by the postnatal laminar environment^15,28^. In contrast, E13-born neurons showed mixed L2/3-like and L5-like features. However, this heterogeneity did not correspond to complete environmental respecification, as E13-born neurons did not uniformly acquire a canonical L5-like identity even when located in deep positions. Rather than indicating broad unrestricted environmental responsiveness, the E13-born outcome may reflect a combination of intrinsic early-born heterogeneity and incomplete or disrupted deep-layer identity acquisition after transplantation^29,37^.

The emergence of a shared, grafting-associated state across all donor groups (Fig. 5a, b) does not undermine this interpretation. Rather, the birthdate-associated difference was superimposed on this common state. The persistence of an E13-versus-E15 distinction despite a shared perturbation argues that birthdate-related programs are intrinsic and robust, as they withstand the dissociation, purification and transplantation procedures that otherwise partially homogenize donor-cell states.

Several mechanisms may account for this asymmetric outcome. The P0 host cortex may favor survival, migration or maturation of IT-like neurons, consistent with the broad L2/3-like bias observed after transplantation^33,38^. Alternatively, the developmental window required for deep-layer identity consolidation may have already narrowed or closed by the time E13-born neurons are transplanted into the postnatal host. In this scenario, early-born neurons may miss temporally restricted embryonic cues, projection-target interactions or circuit-integration windows required for full L5 maturation^39,40^. These possibilities suggest that transplanted neuron identity reflects an interaction between intrinsic birthdate-dependent competence, environmental permissiveness and transplantation-associated perturbation, rather than a simple readout of either birthdate or final laminar position alone.

The use of Patch-seq was important because laminar identity is not captured by a single marker or modality^1,24^. By measuring morphology, electrophysiology and transcriptomic state in the same cells, we found that transplanted neurons could differentiate along several dimensions while remaining only partially aligned with endogenous L2/3 or L5 reference states. The multimodal model validated the asymmetry of this response: E15-born neurons consistently aligned with a superficial/L2/3-like reference state, whereas E13-born neurons occupied a broader mixed space. Thus, the laminar environment alone is insufficient to fully instruct post-mitotic neuron identity, and the degree and direction of differentiation depend on donor birthdate.

This study has several limitations. The design does not fully separate genuine laminar instruction from effects linked to dissociation, FACS purification and transplantation. In addition, we compared transplanted neurons mainly along an L2/3-to-L5 axis, although E13-born populations are intrinsically diverse and include several deep-layer projection-neuron classes^29,37^. Therefore, the mixed E13-born state should not be interpreted simply as failed L5 specification; it may also reflect the broader repertoire of early-born identity programs sampled by this donor population. Finally, radial position is only a proxy for the local environment; future work should examine projections, synaptic inputs and activity-dependent integration. Transplantation into earlier host environments, or into hosts with altered laminar or subtype-specification cues, could further test whether the mixed E13-born outcomes reflect loss of an appropriate temporal environment, intrinsic fate heterogeneity or transplantation-associated disruption.

In summary, our findings show that final laminar position is not sufficient to reprogram the identity of post-mitotic cortical glutamatergic neurons. Instead, donor neurons retain birthdate-dependent differentiation biases, with late-born neurons remaining strongly constrained toward a superficial/L2/3-like state and early-born neurons displaying broader, mixed differentiation outcomes that did not converge on a canonical deep-layer state. These results define an asymmetric window of post-mitotic competence, in which the postnatal cortex can support neuronal integration and maturation but does not substitute for the temporally restricted developmental cues required to consolidate distinct neuron identities.

## Methods

### Animals

Experiments were performed using wild-type CD1 mice from Charles River and CAG::mRFP1 mice (B6.Cg-Tg(CAG-mRFP1)1F1Hadj/J, JAX 005884), previously bred on a CD1 background. Animals were housed in institutional animal facilities under a 12h:12h light/dark cycle with food and water available ad libitum. Timed pregnancies were obtained by overnight mating, and embryonic day 0.5 (E0.5) was defined as the day of vaginal plug detection. Both male and female embryos were used indiscriminately throughout the study. All experimental procedures were approved by the Geneva Cantonal Veterinary Authorities, Switzerland.

### In utero FlashTag labelling

*In utero* FT labelling was performed as previously described. Pregnant CAG::mRFP1 females were anesthetized with isoflurane, the uterine horns were exposed through an abdominal incision and kept humidified with warm 0.9% NaCl. The lateral ventricles of E13.5 or E15.5 embryos were injected with carboxyfluorescein succinimidyl ester (CellTrace CFSE, Life Technologies, C34554) using a picospritzer. The uterine horns were then returned to the abdominal cavity and the abdominal wall was sutured. Embryos were allowed to develop for 48 h before tissue collection.

### Donor cell preparation and FACS purification

Forty-eight hours after FlashTag injection, embryonic brains were collected in ice-cold HBSS (Gibco, 14025092). The dorsal pallium corresponding to the presumptive somatosensory cortex was microdissected under a stereomicroscope. FT^+^/RFP^+^ tissue from one to two litters was pooled and dissociated using the Neural Tissue Dissociation Kit (P) (Miltenyi, 130-092-628), adapted for 1.5 ml tubes. Tissue was incubated in enzyme mix 1 for 5 min at 37°C, followed by enzyme mix 2 for 3 min at 37°C. Cells were mechanically dissociated by pipetting and filtered through a 70 µm cell strainer.

Cells were centrifuged, resuspended in FACS buffer consisting of L15 medium containing 2 mg ml-1 glucose, 0.1% BSA, citrate phosphate dextrose, DNase I and MgCl2, and sorted using a BD FACS Aria II or BD FACSMelody. The 20-30% brightest FT^+^/RFP^+^ cells were collected. Sorted cells were centrifuged and resuspended in HBSS containing 0.1% BSA. The final cell pellet was concentrated to approximately 10 µl with 0.1% Fast Green, yielding approximately 50,000-100,000 cells per µl, and kept on ice until transplantation.

### Neonatal cortical transplantation

Neonatal wild-type CD1 pups at P0.5 were anesthetized with 2% isoflurane (Forene) delivered in a gas mixture containing 30% oxygen and 70% air. A total volume of 0.5–1 µL of concentrated donor-cell suspension was injected into the parenchyma of the presumptive primary somatosensory cortex (S1) using a beveled glass capillary connected to a suction tube.

### EdU labelling

For chronic EdU labelling, osmotic pumps were filled with 10 mg/ml EdU in 1:1 DMSO:water and primed at 37°C for 6-12 h. Pumps were placed in the peritoneal cavity either immediately after FT injection or 24 h later. For pulse labelling, the same EdU solution was injected subcutaneously 1 h before brain collection.

### Tissue collection and immunohistochemistry

From P5 onward, mice were perfused with 4% paraformaldehyde (PFA), and brains were post-fixed overnight at 4°C in 4% PFA before storage in PBS. Coronal sections of 70 µm thickness were cut using a Leica VT100S vibratome. Sections containing transplanted cells were pre-selected under a fluorescence microscope. Free-floating sections were blocked for 1 h at room temperature in PBS containing 5% BSA and 0.25% Triton X-100. Sections were incubated overnight at 4°C with the following primary antibodies, as appropriate: rat anti-RFP (1:500; ChromoTek, 5F8-100), rabbit anti-CUX1 (1:500; Santa Cruz Biotechnology, sc-13024), rat anti-CTIP2/BCL11B (1:500; Abcam, ab18465), rabbit anti-NeuN (1:250; Invitrogen, 14H6L24), and mouse anti-SOX9 (1:500; Thermo Fisher Scientific, AB_2573006). For CTIP2 immunostaining, antigen retrieval was performed in citrate buffer for 20 min at 80°C. Sections were then incubated for 2 h at room temperature with the appropriate secondary antibodies (1:1,000; Thermo Fisher Scientific): goat anti-rat Alexa Fluor 546, donkey anti-rabbit Alexa Fluor 647, donkey anti-rat Alexa Fluor 647, or donkey anti-mouse Alexa Fluor 647. DAPI was added during washing, and sections were mounted with Fluoromount. EdU detection was performed using the Click-iT EdU kit according to the manufacturer’s protocol.

### Imaging and quantification

Images were acquired using Zeiss LSM 700 or LSM 800 confocal microscopes or a Nikon AxR confocal microscope. Quantifications were performed in Fiji/ImageJ. RFP+ cells were manually counted, and marker colocalization was assessed by channel overlay. Unless otherwise stated, only cells located within the cortex were quantified. At least three transplanted brains per experiment were collected and analyzed when applicable. Quantifications were performed using predefined inclusion criteria based on RFP signal, cortical location and marker colocalization. Clumps of cells, particularly those located in the white matter or ventricles, were excluded from analysis. No formal randomization was used. Cell radial position was normalized to cortical depth by dividing the distance from the bottom of the cortex to the cell by the distance from the top of layer 1 to the bottom of layer 6. Laminar position was assigned using DAPI-defined cytoarchitectonic landmarks.

### Single-cell RNA sequencing of donor cells

For pre-transplantation donor-cell profiling, E13.5- and E15.5-born neurons were collected 48 h after FT injection. Cortices from three embryos for the E13.5 time point and six embryos for the E15.5 time point were microdissected and dissociated as described above. Cells were sorted into FACS buffer supplemented with SUPERase-In, RNasin, DNase and magnesium. Approximately 20,000 cells were sorted per time point, and 42.3 µl of cell suspension was loaded onto the 10x Genomics Chromium platform using the 3’ Gene Expression kit. cDNA quality was assessed using Agilent Bioanalyzer and TapeStation systems. Libraries were sequenced at the iGE3 Genomics Platform, University of Geneva. FASTQ files were aligned to the mouse genome GRCm38 using the 10x Genomics Cell Ranger pipeline.

### Patch-seq of transplanted and control neurons

Acute coronal brain slices of 300 µm thickness were prepared from P14-P16 animals. Slices were incubated for 30 min in oxygenated cutting aCSF containing 87 mM NaCl, 2.5 mM KCl, 7 mM MgCl2, 0.5 mM CaCl2, 1.25 mM Na2HPO4, 25 mM NaHCO3, 5 mM glucose and 1 mM kynurenic acid, then gradually transferred to recording aCSF containing 125 mM NaCl, 2.5 mM KCl, 1 mM MgCl2, 2.5 mM CaCl2, 1.25 mM Na2HPO4, 26 mM NaHCO3 and 11 mM glucose. Whole-cell patch-clamp recordings were performed on transplanted RFP+ neurons and neighboring RFP-control neurons located in L2/3 or L5 using 3-4 MOhm pipettes filled with internal solution containing 140 mM KCH3O3S, 2 mM MgCl2, 4 mM NaCl, 5 mM phosphocreatine, 3 mM NaATP, 0.2 mM EGTA, 10 mM HEPES, 0.33 mM GTP and 0.3% biocytin, supplemented with RNase inhibitor. The internal solution was adjusted to pH 7.2. Neurons were recorded in current-clamp mode using 500 ms current steps from 50 to 400 pA, followed by voltage-clamp recordings at −60 mV. For transcriptomic recovery, the nucleus was collected using a nucleated-patch approach and expelled into SMART-Seq v4 3’ DE lysis buffer before storage at −80°C. A total of 120 nuclei were collected from L2/3 and L5 across 20 transplanted animals from five independent transplantation experiments: 11 animals from three E13.5-born transplantation experiments and 9 animals from two E15.5-born transplantation experiments. Of these, 95 nuclei passed quality-control criteria and were retained for analysis. After recordings, slices were incubated to allow biocytin diffusion, fixed overnight in 4% PFA, stained with Alexa Fluor 647-streptavidin and mounted for imaging.

### Patch-seq library preparation and sequencing

cDNA synthesis and amplification were performed using the SMART-Seq v4 3’ DE Kit. Sequencing libraries were prepared with the Nextera XT DNA Library Prep Kit and sequenced on an Illumina HiSeq 4000 using paired-end 50 bp reads, targeting approximately 1 million reads per cell. Reads were aligned to GRCm38 using STAR, and gene counts were generated with HTSeq.

### Morphological reconstruction and analysis

Biocytin-filled neurons were imaged on a Nikon AxR confocal microscope. Neurons were reconstructed in Imaris using the filament tool. Dendritic arbors were resampled into evenly spaced vertices and converted to spherical coordinates using custom MATLAB code. The spatial distribution and orientation of apical and basal dendrites were quantified using probability density functions of radial, azimuthal and elevation components, followed by kernel density estimation and downstream analysis in R.

### Bioinformatic analysis

Single-cell and Patch-seq transcriptomic analyses were performed in R v4.3.2 using Seurat v5.1.0. Visualization was performed with ggplot2 v3.5.2, and doublets were identified using DoubletFinder v2.0.3.

For 10x scRNA-seq datasets, cells were retained if they contained 500-4,000 detected genes, fewer than 5% mitochondrial reads and fewer than 10,000 UMIs. Datasets were normalized, scaled, clustered and visualized using PCA and UMAP. Cell types were annotated manually using established marker genes for apical progenitors, intermediate progenitors, interneurons, migrating neurons and glutamatergic neurons.

Patch-seq datasets were filtered to retain cells with 2,500-10,000 detected genes, fewer than 5% mitochondrial transcripts and fewer than 3,000,000 total counts. Genes detected in fewer than 5% of cells were excluded. Datasets were merged using the intersection of detected genes, and microglia-associated genes were regressed out before downstream analysis.

To quantify the extent to which each molecular, morphological, or electrophysiological modality encoded cortical layer identity, linear support vector machine (SVM) classifiers were trained using control cells only. Separate models were fitted for the Morphology, Electrophysiology, Transcriptomic, and Multimodal datasets with the objective of predicting cortical layers L2/3 and L5.

Models were implemented using the tidymodels framework in R. For electrophysiological and morphological variables, preprocessing recipes consisted of standardization and mean imputation. RNA features were provided to the model as the log-normalized counts without additional preprocessing. The classifier was a linear SVM implemented through the LiblineaR engine, with the regularization cost parameter set to 0.1.

Classifier performance was assessed using leave-one-out cross-validation (LOOCV) on the control-cell dataset. A global model, using all control cells, was trained and used to predict the transplanted cells. The resulting decision value therefore represented the position of each cell along a layer-classification axis learned exclusively from control cells. Classification performance was compared using overall accuracy and the area under the receiver operating characteristic curve (AUC).

To identify features contributing most strongly to layer classification, the fitted coefficients of each linear SVM were extracted.

Variance partitioning was performed on transplanted neurons only and separately for each morphological, electrophysiological, transcriptomic and multimodal identity measure using linear mixed-effects models fitted with lme4 and lmerTest. Donor birthdate, final laminar position and their interaction were included as fixed effects, with mouse included as a random intercept. The unique contribution of each fixed effect was quantified as semi-partial R² using partR2 and expressed as a percentage of the total variance; the remaining variance, defined as 100% minus the summed semi-partial R² contributions, was classified as unexplained.

### Differential expression, heatmaps and gene ontology

Differential expression analysis was performed with Seurat using two-sided Wilcoxon rank-sum tests. L2/3- and L5-specific genes were identified from control Patch-seq neurons using thresholds of Bonferroni-adjusted P < 0.01, minimum expression in 40% of cells and average log2 fold change > 0.1. Average expression values were computed per experimental group and centered relative to control L2/3 and L5 neurons. Genes were categorized as conserved, lost or induced relative to controls and visualized by hierarchical clustering using Ward.D2 linkage. Gene ontology enrichment was performed on gene lists derived from differential expression and heatmap clustering. Over-representation analysis was performed using fgsea, and enrichment outputs were ranked by adjusted P value and visualized as heatmaps. Significant pathways were defined at Benjamini–Hochberg-adjusted P < 0.05.

### Statistics

Unless otherwise stated, statistical analyses were performed in R, and all statistical tests were two-sided. Normality was assessed using the Shapiro–Wilk test. For independent pairwise comparisons, two-sided Welch’s t-tests were used for normally distributed data, whereas two-sided Wilcoxon rank-sum tests were used for non-normally distributed data. For paired comparisons, two-sided paired t-tests or Wilcoxon signed-rank tests were used, as appropriate. Electrophysiological and morphological features were analyzed using two-way linear mixed-effects models including experimental condition, final laminar position and their interaction as fixed effects, with mouse included as a random intercept. Significant main effects or interactions were followed by post-hoc pairwise comparisons of estimated marginal means. Unless otherwise stated, multiple-testing correction was performed using the Benjamini–Hochberg procedure, and significance was defined as an adjusted P value < 0.05. Data are presented as mean ± SEM unless otherwise stated, and sample sizes indicate independent biological replicates or individual neurons, as specified in the figure legends.

### Software and packages

Transcriptomic analyses were performed in R v4.3.2 using Seurat v5.1.0. Data visualization was carried out using ggplot2 v3.5.2, and doublets were identified using DoubletFinder v2.0.3. Morphological analyses were performed using Imaris v8.1.2 for neuronal reconstruction and MATLAB R2022b for custom processing of dendritic morphology. The 10x Genomics scRNA-seq data were aligned to the GRCm38 mouse reference genome using Cell Ranger v7.2. SMART-seq Patch-seq reads were aligned to GRCm38 using STAR v2.7.10a, and gene counts were generated using HTSeq v2.0.2. Additional R packages used for analysis included tidymodels v1.4.0, clusterProfiler, fgsea, LiblineaR, lme4, lmerTest, emmeans, partR2, bmrm v4.1, and stats v4.3.2.

## Acknowledgements

We thank the iGE3 Genomics Platform, Bioimaging, and FACS Facility at the University of Geneva, the Human Cellular Neuroscience Platform at Campus Biotech; A. Benoit for technical assistance; the entire Jabaudon laboratory for their thoughtful feedback on the manuscript and for their constructive contributions throughout the project. The Jabaudon laboratory is supported by the Swiss National Science Foundation, the European Research Council and the NeuroNA foundation. R.B. was supported by the Swiss National Science Foundation (Ambizione grant: PZ00P3_201995). S.R.-P. was supported by an EMBO Long-Term Fellowship (ALTF 349-2020).

## Author contributions

E.B. and D.J. conceived the project and designed experiments. E.B. performed transplantation experiments, histological analyses and immunohistochemistry with input from R.B. and D.J. S.F. contributed to Patch-seq experiments, S.R.-P. to 10x scRNA-seq experiments, and C.F. to morphology and Smart-seq analysis pipelines. E.B., J.P., R.B. and D.J. analyzed data and interpreted results. R.B. and D.J. supervised the project. R.B. shaped the conceptual framework, integrated the multimodal results into the final story, and wrote the manuscript with input from E.B., D.J. and all authors.

---

**Supplementary Fig. 1.**
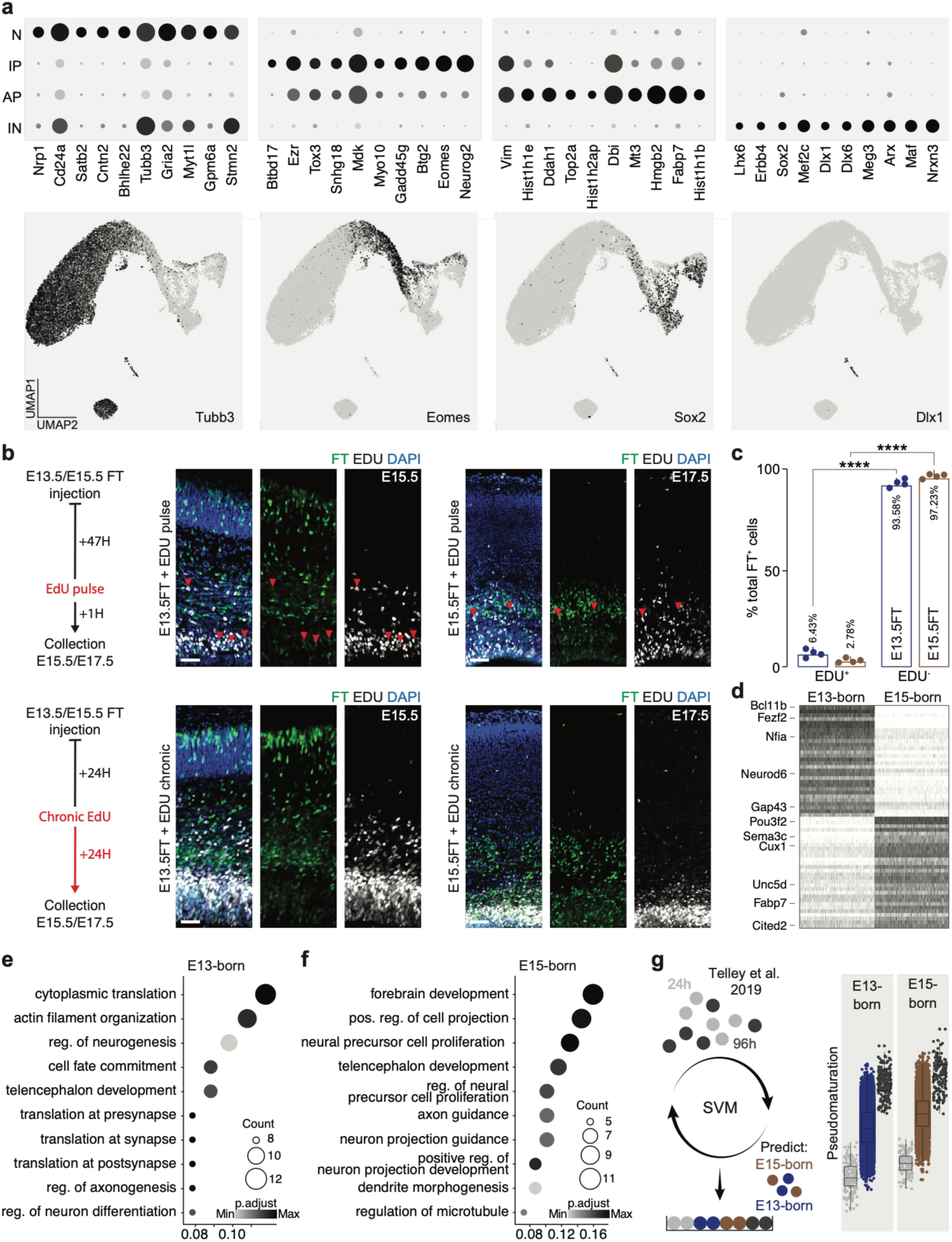
Donor populations are enriched for post-mitotic neurons and show birthdate-dependent transcriptional differences. **(a)** Expression of the top ten marker genes for each cell type. **(b)** EdU pulse and chronic EdU labelling designs, with representative images before donor collection. **(c)** Quantification of EdU-positive and EdU-negative FT-positive donor cells after EdU pulse labelling. n = 4 mice. Two-sided paired t-tests, with Benjamini–Hochberg correction. **(d)** Heat map of differentially expressed genes between E13-born and E15-born donor neurons before transplantation. E13-born neurons preferentially express DL-associated genes, whereas E15-born neurons preferentially express SL-associated genes. **(e, f)** Gene ontology enrichment for genes enriched in E13-born (e) and E15-born (f) donor neurons. Dot size indicates gene count, and color intensity indicates adjusted p value. **(g)** Pseudomaturation SVM classifier trained on published FT-labelled neurons collected 24 h and 96 h after labelling, then applied to 48 h donor neurons to assign each cell a maturation score. ****p < 0.0001. Bars show mean +/− SEM.

**Supplementary Fig. 2.**
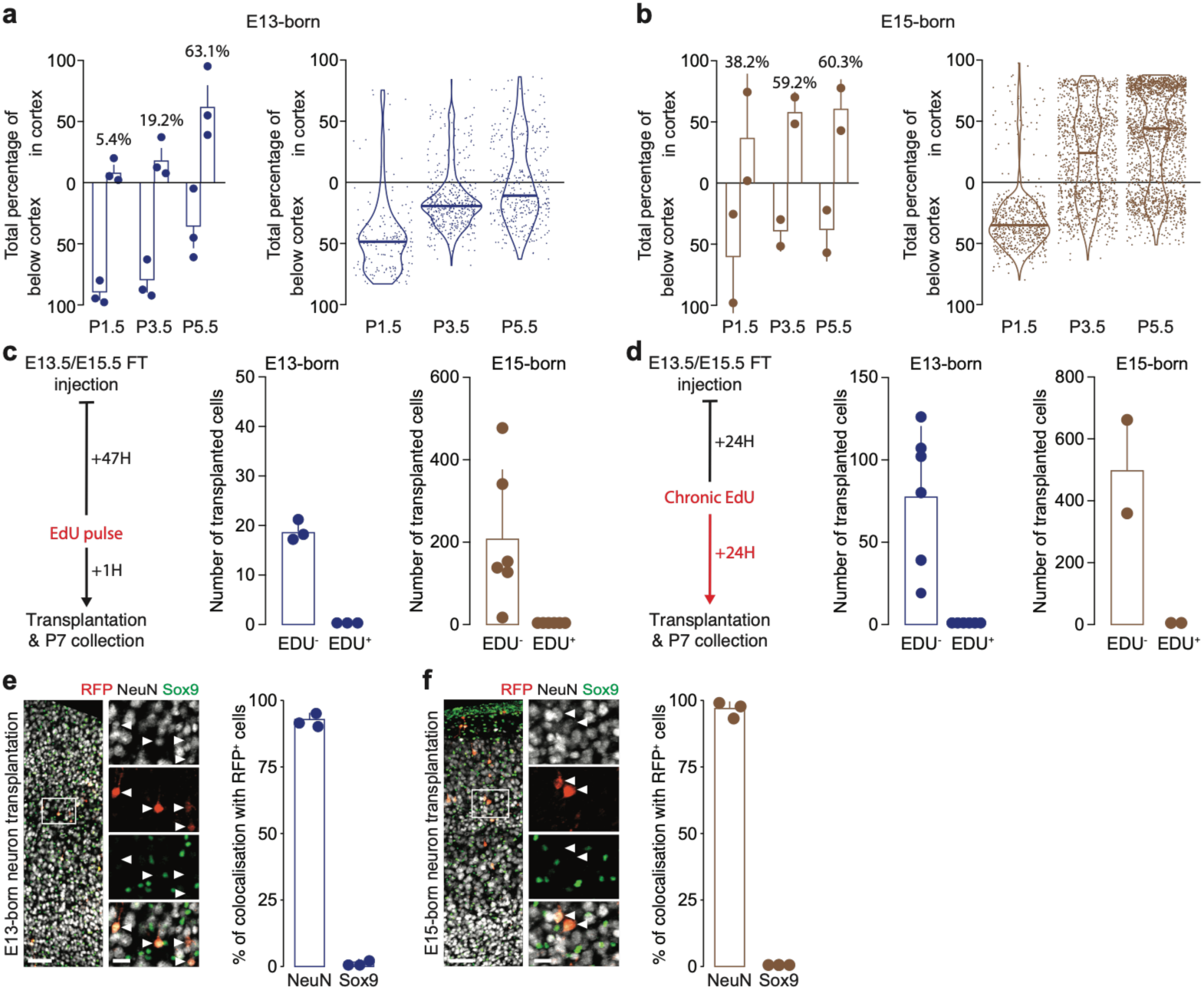
Validation of transplanted-cell position, cell-cycle status, and neuronal identity. **(a, b)** Quantification of E13-born (a) and E15-born (b) transplanted cells located within or below the cortex at early postnatal stages. Bar plots show the percentage of cells found within or below the cortex, while violin plots show the normalized radial distribution of individual cells. n = 3 mice. **(c, d)** EdU pulse (c) and chronic EdU (d) paradigms, together with quantification of EdU-negative and EdU-positive transplanted cells at P7. No integrated transplanted neurons were EdU-positive, indicating that cortical integration did not result from donor cells cycling shortly before transplantation. For the EdU pulse experiment, n = 3 transplanted mice for E13-born cells and n = 6 for E15-born cells; for the chronic EdU experiment, n = 3 transplanted mice for E13-born cells and n = 6 for E15-born cells. **(e, f)** Representative images and quantification showing that RFP-positive E13-born (e) and E15-born (f) transplanted cells co-localize with NeuN and show little to no co-localization with Sox9. n = 3 mice. Bars show mean +/− SEM. Scale bars, 100 μm; insets, 15 μm.

**Supplementary Fig. 3.**
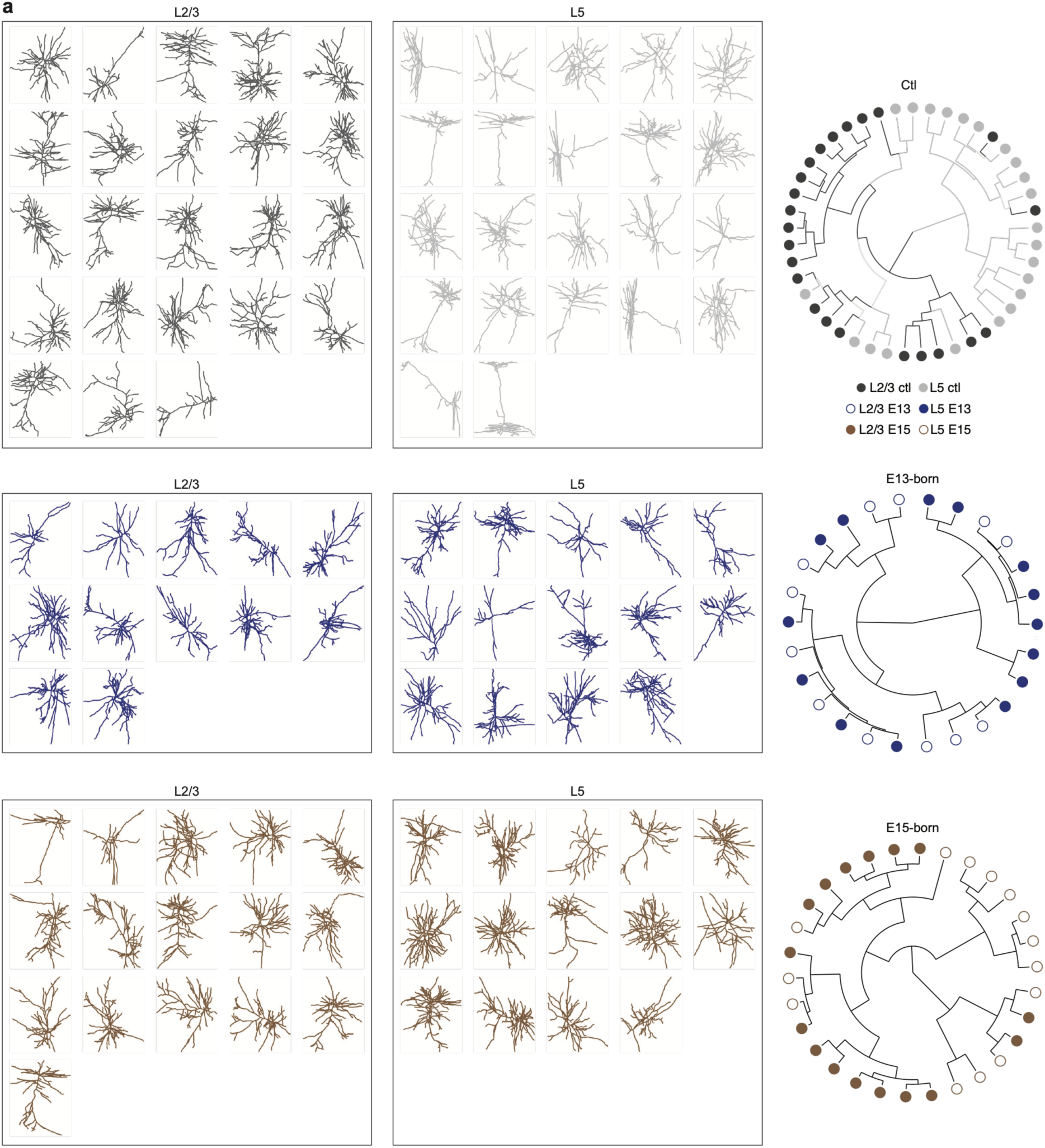
Hierarchical clustering of reconstructed neuronal morphologies. **(a)** Reconstructed morphologies of all control, E13-born and E15-born neurons analyzed in this study and located in L5 or L2/3 (left), alongside circular hierarchical clustering (right) of manually reconstructed neurons based on discriminative morphological features (Supplementary Fig. 4a). Reconstructions are colored by group and final laminar position, illustrating the morphological diversity of control and transplanted neurons.

**Supplementary Fig. 4.**
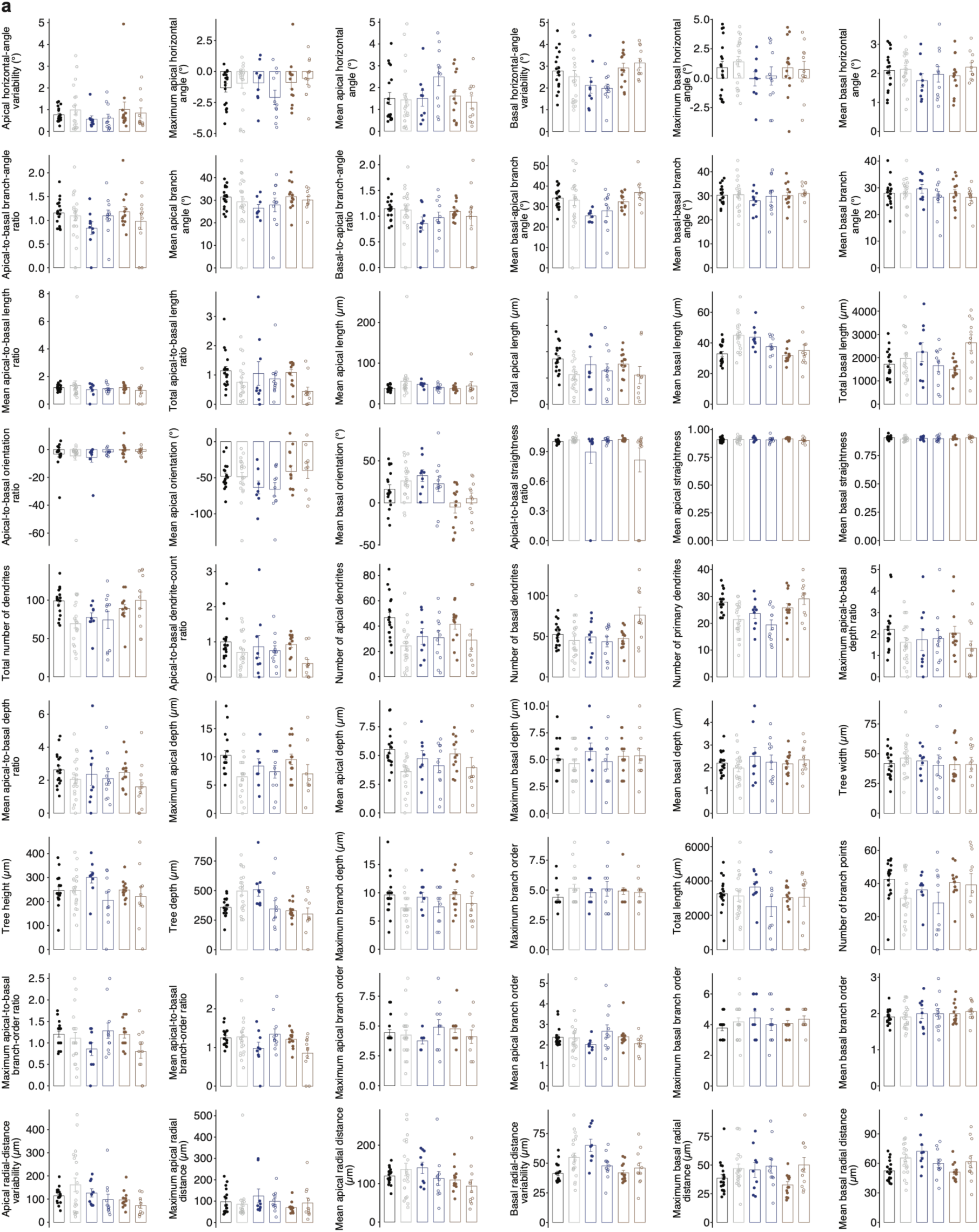
Individual morphological features across groups. **(a)** Quantification of morphological parameters across control L2/3, control L5, E13-born L2/3, E13-born L5, E15-born L2/3, and E15-born L5 neurons. n = 21 neurons for L2/3 ctl, 22 for L5 ctl, 14 for L2/3 E13, 13 for L5 E13, 13 for L2/3 E15, and 12 for L5 E15. Bars show mean +/− SEM.

